# Pcbp1 constrains Oct4 expression in the context of pluripotency

**DOI:** 10.1101/2024.09.07.611681

**Authors:** E. I. Bakhmet, A. S. Zinovyeva, A. A. Kuzmin, D. V. Smirnova, M. N. Gordeev, E. E. Petrenko, N. D. Aksenov, A. N. Tomilin

## Abstract

Oct4 is a commonly known marker of pluripotent stem cells as well as one of the key factors required for pluripotency induction. Its gene (*Pou5f1*) is subject to complicated regulation through distal and proximal enhancers. Noteworthy, this protein also plays an important role in primitive endoderm (PrE) specification, though the mechanisms driving its expression during this process are still unknown. Here we show that KH-domain protein Pcbp1 occupies poly(C)- sites of the *Pou5f1* enhancers, but *Pcbp1* knockout does not affect the Oct4 expression level in ESCs. On the contrary, Pcbp1 is essential for timely Oct4 downregulation upon differentiation signals. Indeed, Pcbp1 loss results in high rate of spontaneous differentiation of ESCs to PrE, and this effect is enhanced upon retinoic acid treatment. This phenotype can be explained by residual Oct4 expression, as Oct4 depletion rescues normal differentiation. Overall, our results point to Pcbp1 is a transcriptional regulator of *Pou5f1*, purported to synchronize Oct4 expression decline with the pluripotency network shutdown during differentiation. Oct4 being outside of this network loss its functions as factor of pluripotency and acts as PrE specifier.

## Introduction

Oct4 is a pioneer transcription factor which is mainly known for its functions in the induction and maintenance of pluripotency, as well as in survival of primordial germ cells (PGCs) [1-5]. On the other hand, a body of evidence suggests instructive Oct4 functions in differentiation of pluripotent stem cells. It has been shown that during germ layers specification, Oct4 and Sox2 promote mesoendoderm and neuroectoderm formation, respectively [6-9]. Oct4 expression in early embryogenesis is also essential for primitive endoderm (PrE) development [10-13]. While Oct4-Sox2 cooperativity is important for pluripotency maintenance [14], Oct4 partnership with Sox17 is critical for PrE differentiation [15, 16]. *Pou5f1* gene expression is regulated through its distal enhancer (DE), proximal enhancer (PE), and promoter [17-21]. It is known that the DE is active in pluripotent cells of epiblast before implantation and in their cultured counterparts – naïve ESCs, and that the PE drives Oct4 expression in post-implantation epiblast cells and in its cultured counterparts – the epiblast stem cells (EpiSCs), while both enhancers function in PGCs [17, 21]. Though there is a significant amount of information concerning proteins that bind these two regulatory elements [18, 22-24], many gaps in understanding the mechanisms by which they control *Pou5f1* gene transcription at different timepoints of early development remain to be fulfilled.

The family of KH-domain poly(rC)-binding proteins includes five members – hnRNP-K and Pcbp1-4. These factors are characterized by their ability to bind DNA/RNA (T/U)CCC-motifs [25-29]. The broad spectrum of functions for these proteins including transcriptional regulation [30-33], mRNA-processing [34-36], splicing [37-40], translation [29, 41-43], and iron metabolism [44-46] were described. By taking an affinity purification approach we have previously identified hnRNP-K, Pcbp1, and Pcbp2 as proteins binding the poly(C)-motif 2A of the *Pou5f1* DE *in vitro* [47, 48]. HnRNP-K has been found essential for viability of both ESCs and differentiated cells, but, surprisingly, not engaged in activation, maintenance, or downregulation of *Pou5f1* gene in these cells, even though it occupies *in vivo* the poly(C)-motifs 2A and 1A of the DE and PE, respectively [48]. Another identified 2A-binding protein, Pcbp1, is also known to function during embryogenesis. This protein participates in transcription silencing in murine oocytes [49], while a recent study pointed to Pcbp1/Pcbp2 function in the promotion of trophectodermal lineage choice during early embryogenesis via *Carm1* pre-RNA exon-skipping splicing [37]. Despite high Pcbp1/Pcbp2 structural and functional similarity, their roles in further development are different. *Pcbp1* knockout leads to peri-implantation lethality through some unknown mechanisms, while Pcbp2 ablation leads to broad cardiovascular and erythropoiesis abnormalities at further developmental timepoints [50, 51].

We have also shown that Pcbp1-deficient ESCs (hereafter referred to as KO), generated using the CRISPR/Cas9 tool, express normal levels of Oct4, show self-renewal capacity, however, are prone to spontaneous differentiation [52]. Therefore, we have questioned (1) whether Pcbp1 binds *Pou5f1* the abovementioned regulatory elements in ESCs, (2) to what type of spontaneous differentiation leads Pcbp1 deficiency, and (3) if these two are linked to each other. In this study we show that Pcbp1 binds the DE and PE of *Pou5f1* gene in ESCs and is needed for timely Oct4 down-regulation during ESC differentiation. Overall, our study confirms Pcbp1 functions in transcriptional regulation of the *Pou5f1* gene and clarifies the mechanisms driving Oct4-mediated PrE differentiation.

## Methods

### Cell culture

Unless specified, all reagents for mouse ESC culturing were purchased from Gibco (ThermoFisher Scientific). ESCs were grown on adhesive plastic (Eppendorf, TPP) pre-coated with 0.1% gelatin (Merck) under standard culture conditions (5% CO_2,_ 37ºC) in the Knockout-DMEM medium supplemented with 15% of fetal bovine serum (HyClone), 1x penicillin/streptomycin, 2 mM of L-Glutamine, 1x non-essential amino acids, 50 μM β-mercapthoethanol, and 500 U/ml of bacterially expressed human LIF made in in-house. The cell culture condition was designated as the SL. When mentioned, 1 μM PD0325901 (Axon), 3 μM CHIR99021 (Axon) were included. Passaging was performed using 0.05% Trypsin-EDTA. For all-trans retinoic acid (RA)-mediated differentiation, ESCs were collected, washed with PBS, and seeded onto 0.1% gelatin-coated plastic dishes at a various density, depending on experiment (usually 10^4^ cells/cm^2^ for three-day differentiation) in medium supplemented with 1 μM RA (Merck) and deprived of LIF for indicated periods of time. For experiments with preliminary Oct4 depletion in “floxed” ESCs, 0.8 μg/ml of 4-hydroxytamoxifen (4-OHT, Merck) was added to the culture medium 24 hours before RA-driven differentiation. All cell lines were routinely checked for mycoplasma contamination and all were negative.

### DNA constructs

The plasmids *pRosa26-TRE-CAG-Frt(Ert2CreErt2-STOP)Frt-tdTomato-PGKneo* harboring *Cre* gene flanked by homology arms to *Rosa26* locus, as well as the plasmid *pX330-U6-Chimeric_BB-CBh-hSpCas9* harboring *EGFP* gene and *Rosa26*-targeting gRNA were reported previously [53]. The *dtTomato* gene in the first construct is not expressed due to the stop codon and polyadenylation signal after Cre-coding sequence. For *T2A-EGFP* integration into the *Pou5f1* locus, the plasmid *phCas9D10A-Pou5f1* harboring Nickase and *Pou5f1-*targeting gRNA (5’-GTCTCCCATGCATTCAAACTG -3’), as well as the plasmid *pOct4-T2A-EGFP* harboring *T2A-EGFP* sequence flanked by 4342-bp left and 2754-bp right homology arms were used for *T2A-EGFP* sequence insertion just upstream of the *Pou5f1* stop codon.

### Mutant ESCs

The *Oct4*^*T2A-EGFP*^ Scr and KO reporter ESC lines were generated from the Tg2a ESCs (Bay Genomics) using the described above DNA constructs. The *Pcbp1*^− */*−^ (KO) and isogenic scrambled control ESC (Scr) derivation was reported elsewhere [52]. The *Pou5f1*^*flox/flox*^;*Rosa26*^*Ert2CreErt2*^ ES cell line was derived from the *Pou5f1*^*flox/flox*^ ESCs [5], by inserting the *Ert2-Cre-Ert2* into the *Rosa26* locus, using previously designed and described above DNA constructs [53]. *Pcpb1* gene knockout in the “floxed” ESCs was generated as described previously [52]. “Floxed” ESC clone 2 was used in subsequent experiments.

### Chromatin immunoprecipitation

ChIP assay was performed as described elsewhere [48]. In brief, cells were fixed directly on 6-cm culture dishes with 1% formaldehyde, quenched by 0.125 M Glycine, washed with PBS, scrapped, and resuspended in Sonication buffer with protease inhibitor cocktail (PIC, Roche). Then, chromatin was fragmented using the sonicator CL-18 (ThermoFisher Scientific) for 10 seconds with 60% amplitude 10 times with 60-sec cooling on ice after each cycle. Afterwards, immunoprecipitation was performed as follows: chromatin from 10^6^ cells was incubated with 5 µg pre-immune rabbit IgG or IgG to Pcbp1, conjugated with MabSelect sure Sepharose (Cytiva). Precipitated IgG-protein-DNA complexes were eluted, and crosslinking reversal was performed by incubation with proteinase K, NaCl, and EDTA at 55ºC for 10 hrs and 65ºC for 5 hrs. DNA was purified by phenol:chloroform extraction and dissolved in 30 µl of TE buffer; 2 µl of DNA were taken for qPCR.

### Cell staining

Cells were fixed with 4% paraformaldehyde for 10 min directly on culture plates, washed with PBS plus 0.1% Tween (PBST), permeabilized by 0.1% Triton X-100 for 15 min, and blocked with 3% BSA-PBST for 30 min. Incubation with the primary antibodies, diluted in 3% BSA-PBST, was performed for 2 hrs at room temperature or overnight at +4°C. Cells were washed 5-6 times with PBST and incubated with secondary fluorescent antibodies diluted in 3% BSA-PBST for 1 hr at room temperature. Afterwards, cells were washed 3 times with PBST, incubated with DAPI for 2 min, washed, and stored in PBS with sodium azide. Imaging was performed by EVOS fl Auto (ThermoFisher Scientific). Alkaline phosphatase staining was performed with Alkaline Phosphatase Kit (Merck) in accordance with manufacturer recommendations. Immunofluorescent staining for the flow cytometry was performed with True-Nuclear™ Transcription Factor Buffer Set (BioLegend) according to manufacturer recommendations.

### Quantitative mRNA analysis

Total RNA from the cells cultured in 6-well plates was isolated using the ExtractRNA kit (Evrogen). Reverse transcription with 1 µg of the RNA was performed using the MMLV RT kit (Evrogen). RT-PCR was performed with 2.5 µL of cDNA and SYBRGreen PCR Mastermix (Evrogen), using the LightCycler® 96 System (Roche). The thermocycling conditions were as follows: 95°C for 5 min, followed by 45 cycles of 95°C for 15 sec and 60°C for 1 min. A final heating step from 65°C to 95°C was performed to obtain the melting curves of the final PCR products. Changes in target gene expression levels were calculated as fold differences using the comparative ddCT method. The mRNA levels were normalized to glyceraldehyde 3-phosphate dehydrogenase (Gapdh) mRNA. Statistical analysis was performed using GraphPad Prism software (version 8.0.1, GraphPad Software).

### Western blot

For Western blot analysis, cells were harvested from 6-well plates by trypsinization and resuspended in PBS. Aliquot (1/10) of this mixture was used for protein concentration measurement by Pierce™ BCA Protein Assay Kit (ThermoFisher Scientific). Then 2x Laemmli buffer with β-mercaptoethanol was added and probes were boiled at 100ºC for 5 min. For electrophoresis, 10-20 µg of the total protein from each probe were taken, and separation was performed at 80 V in 10% acrylamide gel. Nitrocellulose membrane (Bio-Rad) was used to transfer proteins from gel, which then was blocked by 5% non-fat Milk diluted in PBST.

Incubation with primary antibodies was performed in 3% BSA-PBST for 2 hrs at room temperature. After 3-4 washes in PBST membranes were incubated with secondary antibodies diluted in 1% Milk in PBST for 1 hr, then washed three times by PBST. Chemiluminescence was visualized using the SuperSignal™ West Pico PLUS Chemiluminescent Substrate (ThermoFisher Scientific) and ChemiDoc™ Touch Imaging System (Bio-Rad).

### Cell cycle analysis

Cells were harvested by trypsinization, washed with PBS, and resuspended in PBS supplemented with 0.025 % saponin, 50 µg/mL propidium iodide (PI), and 0.25 mg/mL RNAse A for 40 minutes at RT. The cells were then analyzed by flow cytometry, and cell cycle distribution was assessed using ModFit LT 3.0 software (Verity Software House, Lexington, MA, USA).

### Flow cytometry and FACS

Flow cytometry was performed using the CytoFLEX benchtop flow cytometer (Beckman Coulter) with lasers wavelengths 488 nm for EGFP detection (FITC) and 638 nm for Alexa 647-conjugated secondary antibody detection (APC). Fluorescent-activated cell sorting was performed using the S3e Cell Sorter (Bio-Rad) in decontaminated conditions.

### Statistical analysis

Statistical analysis was performed using GraphPad Prism. Unless specified otherwise, data were analyzed using an unpaired Student’s t-test. Values are expressed as means ± SD. “N” designated biological replicates, and “n” designated technical replicates. The significance levels were as follows: ns, not significant; *p < 0.05; **p < 0.01; ***p < 0.001; ****p < 0.0001.

**Table 1.**
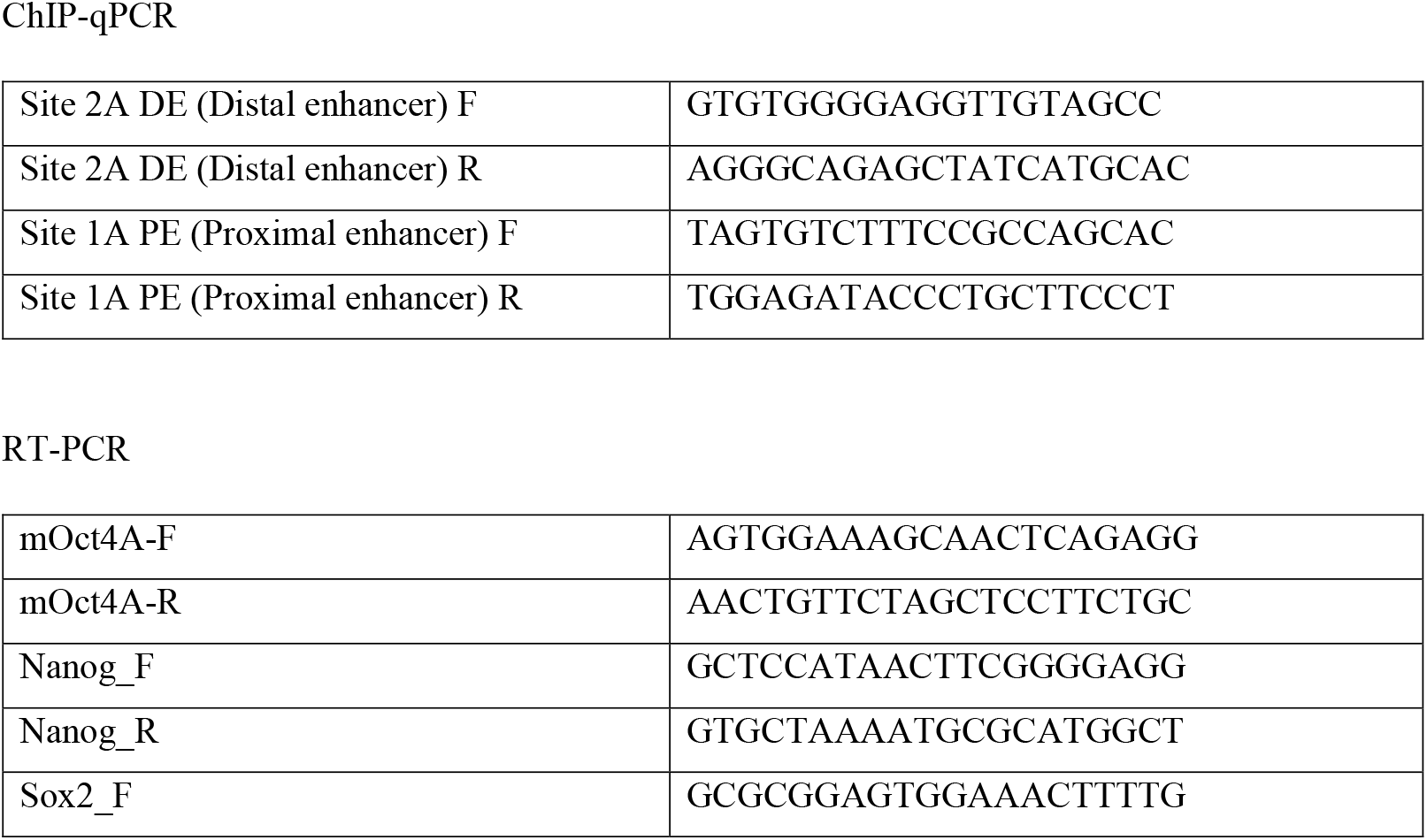
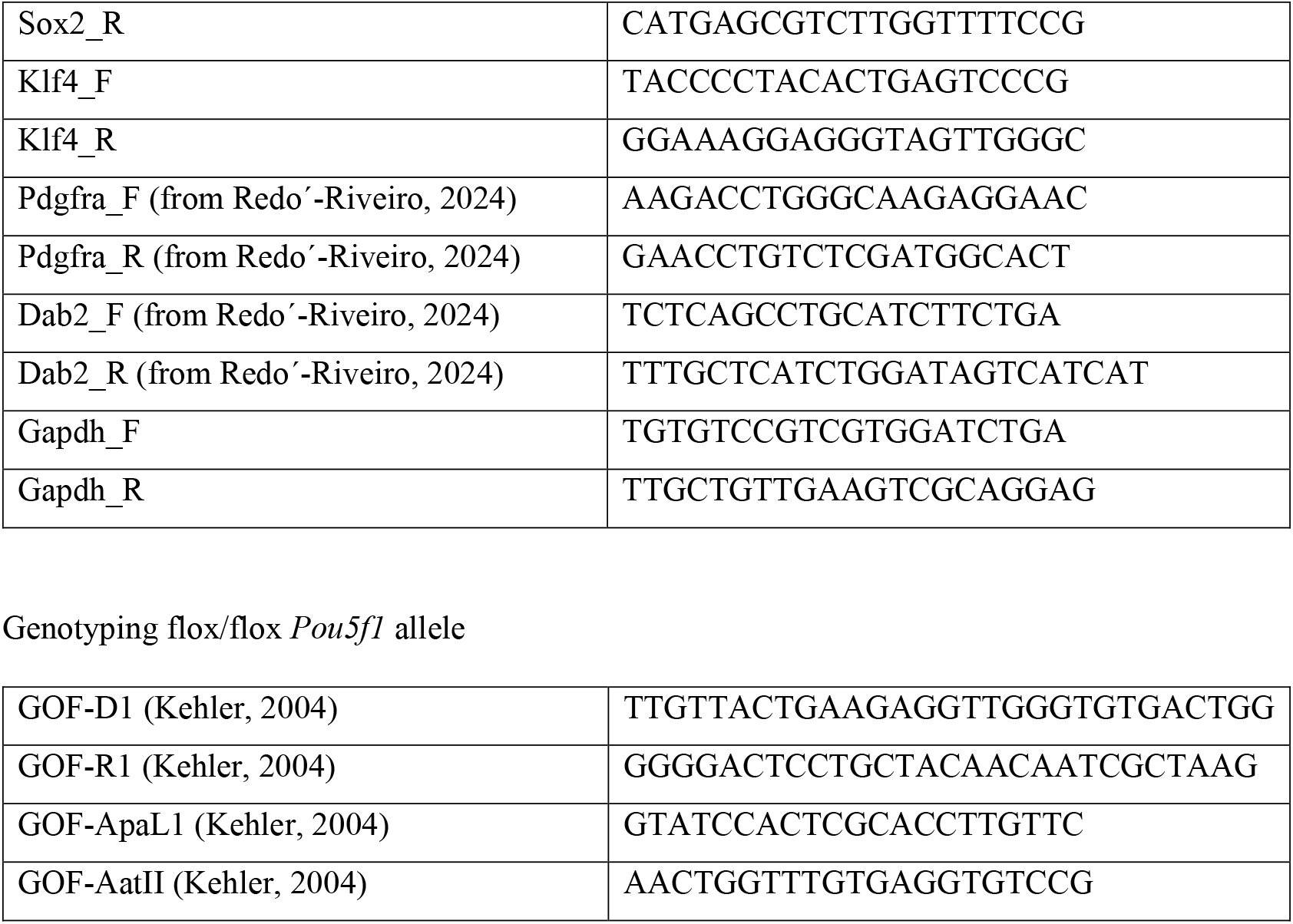
Primers used in this study.

**Table 2.** Antibodies used in this study.

| Target protein | Cat. No. | Manufacturer |
| --- | --- | --- |
| Pcbp1 | ab74793 | Abcam |
| Oct4 | sc-5279 (C-10) | Santa Cruz |
| Nanog | A300-397 | Bethyl |
| Gata6 | AF1700 | RD systems |
| Sox17 | AF1924 | RD systems |
| Gapdh | 2118 | Cell Signaling |
| β-actin | 643802 | BioLegend |

Secondary fluorescent and HRP-conjugated antibodies were purchased from Jackson ImmunoResearch.

## Results

### Pcbp1 binds *Pou5f1* enhancers in ESCs

Considering the fact that Oct4 expression is not affected in undifferentiated KO ESCs [52] we first asked whether Pcbp1 occupies in ESCs the poly(C)-sites 2A and 1A of, respectively, the DE and PE of the *Pou5f1* gene. The cells were cultured in the presence of serum and LIF (referred hereafter to as the SL condition), known to potentiate activity of the both enhancers [17, 21]. We have also analyzed differentiated cells which are derived from ESCs by means of all-trans retinoic acid (RA) treatment (along with LIF withdrawal) and characterized by a rapid Oct4 down-regulation (Fig. 1A) [20]. Following three days of exposure to RA, cells showed complete pluripotency shutdown, as evidenced by the loss of Oct4 and Nanog expression (Suppl. Fig. 1A, B). As evidenced by ChIP-qPCR, Pcbp1 occupies in ESCs both the 2A and 1A sites of the *Pou5f1* DE and PE, respectively, and, albeit the protein is expressed in differentiated cells at approximately same level (Suppl. Fig. 1A), it shows about 5-fold occupancy reduction in the latter cells (Fig. 1B). These observations indicate that there are some mechanisms restricting Pcbp1 binding in differentiated cells. Overall, our data shows that Pcbp1 occupies the two well-known *Pou5f1 cis*-elements *in vivo* and thus, might be involved in transcriptional regulation of this gene.

**Figure 1.**
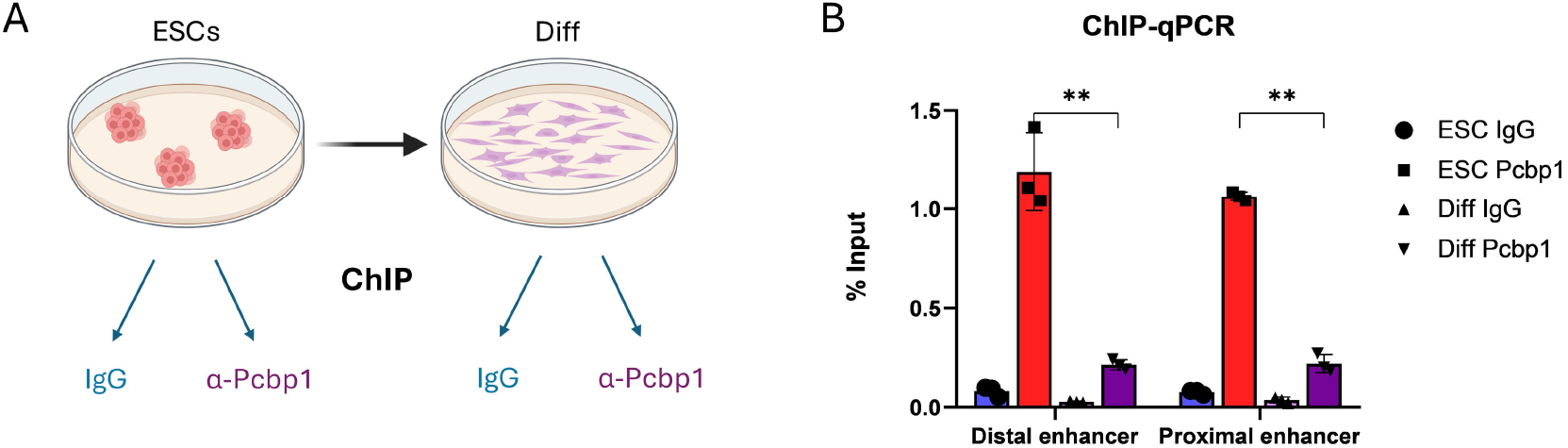
Pcbpl occupies *Pou5f1-* regulatory regions in ESCs. (A) Schematic representation of ChIP analysis of ESCs cultured in medium containing serum and LIF (SL) and differentiated via RA exposure for three days; (B) ChIP-qPCR analysis of Pcbpl binding to the sites 2A and IA of the *Pou5fl* distal (DE) and proximal enhancers (PE), respectively (n=3).

### Pcbp1-deficient ESCs are prone to spontaneous differentiation into PrE

Previously established Pcbp1-deficient ESCs, being capable of giving rise to all three germ layer derivatives within teratomas, showed clear signs of spontaneous differentiation in culture [52]. Noticeably, these cultures contain cells which possess cobblestone morphology and are negative for alkaline phosphatase – the known feature of primitive endoderm (PrE) on adhesive substrates (Fig. 2A). Indeed, Western blot analysis showed the presence of Gata6 and Sox17 proteins in cell lysates of the KO ESCs (Fig. 2B), while RT-PCR revealed the expression of Pdgfra and Dab2 mRNAs which are distinctive markers of PrE (Suppl. Fig. 2A).

**Figure 2.**
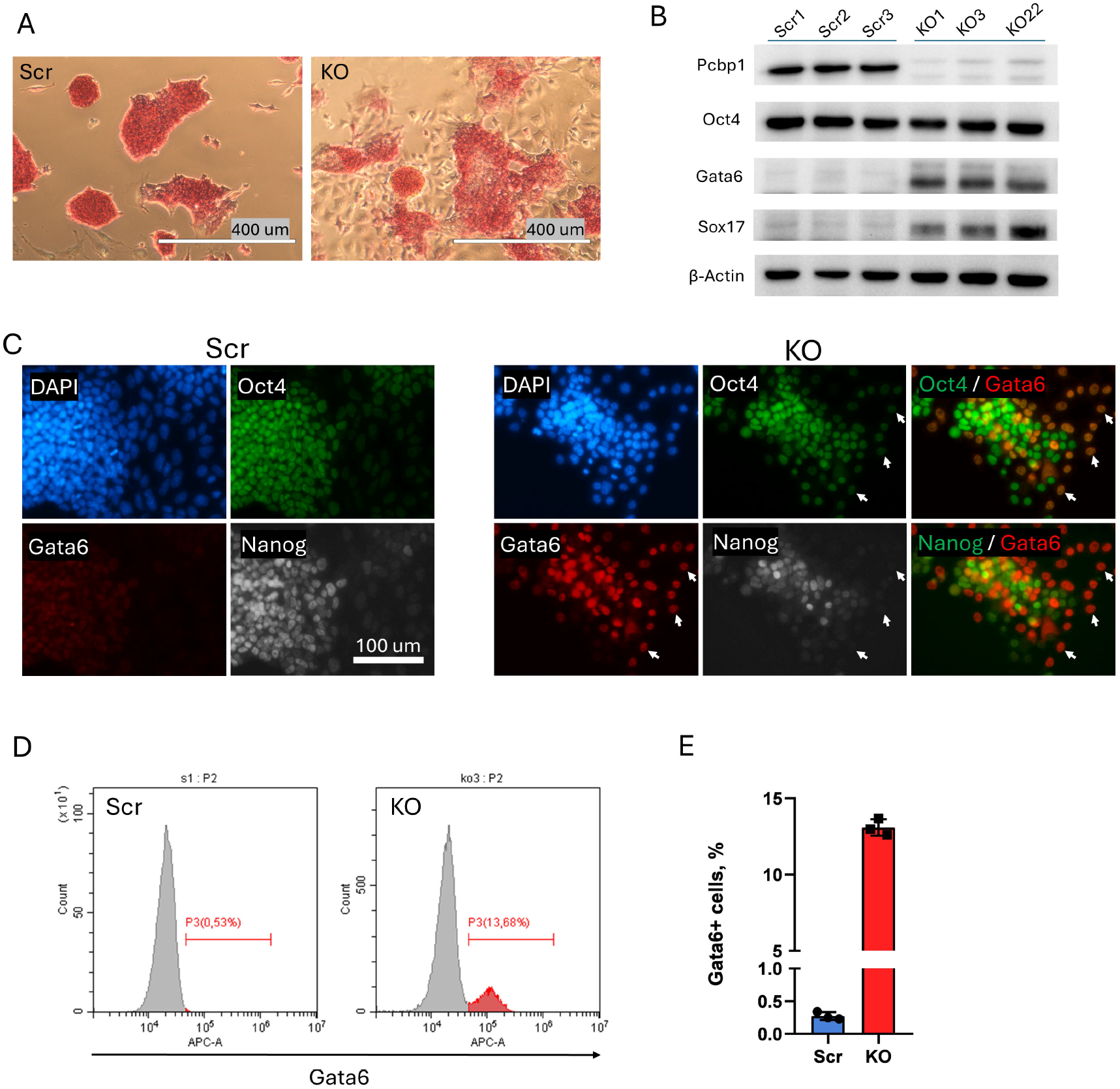
Pcbpl function loss in ESCs promotes spontaneous differentiation to PrE. (A) Alkaline phosphatase staining of the control (Ser) and Pcbp I -deficient (KO) ESCs cultured under the SL conditions. (B) Western blot showing the expression of Pcbpl, Oct4, and the PrE markers Gata6 and Sox 1 7 in independently derived ESC clones of indicated genotype. (C) Immunofluorescence staining of Ser and KO cells using Oct4, Nanog, and Gata6 antibodies; white arrows on the right panel indicate Oct4+Gata6+Nanog-PrE cells. (D) Typical flow cytometry estimation of Gata6+ cell percentage in Ser (left) and KO ESCs (right). (E) Mean Gata6+ cell percentage across independently derived Ser and KO ESC lines (N=3 for each).

Immunocytochemical staining showed rare Gata6^+^ cells within Scr but numerous Gata6^+^ cells within KO ESC populations (Fig. 2C). Noteworthy, these cells were negative for Nanog and positive for Oct4 (Fig. 2C, right panel, white arrows). This observation agrees with the known Nanog–Gata6 antagonism [54-56], as well as with the long-standing view of Oct4 function in PrE specification [10, 13]. Finally, Gata6 staining with consequent flow cytometry analysis revealed that nearly 13% of KO cells are represented by PrE (Fig. 2D, E).

Further characterization of the KO ESCs showed a reduced, as compared to the Scr ESCs, proliferation rate (Suppl. Fig. 2B), though there was no obvious re-distribution between cell cycle phases (Suppl. Fig. 2C). To rule out the possibility that the PrE differentiation of the KO ESCs is triggered by the affected proliferation rate, we cultured the Scr ESCs in a medium containing incrementally decreased concentrations of fetal bovine serum. This predictably resulted in correspondingly reduced proliferation rates (Suppl. Fig. 2D) but did not promote PrE differentiation (Suppl. Fig. 2E). Thus, it could be concluded that Pcbp1-deficiency endorses spontaneous differentiation of ESCs to PrE cells via some mechanisms not related to cell cycle control. Also, our data indicate Pcbp1 involvement into regulation of some basic cellular processes because its loss leads to an increase of the overall cell cycle length.

### Pcbp1 is needed for Oct4 decline during ESC differentiation

Our results clearly show that Pcbp1 occupies *Pou5f1* regulatory regions, however, its loss does not alter Oct4 expression in ESCs under the SL conditions (Fig. 2B, C). We next asked if Pcbp1 is needed for timely Oct4 down-regulation during differentiation of ESCs, such as this induced by RA (along with LIF withdrawal). As expected, the treatment leads to a rapid Oct4 down-regulation in control Scr cells (Fig. 3A, left panel), whereas in the KO cells, Oct4 expression persists in a significant subpopulation with most cells within this subpopulation additionally expressing Gata6 (Fig. 3A, right panel). However, further differentiation of the KO cells is accompanied by a loss of the Oct4^+^/Gata6^+^ sub-population, which could be due to incompatible with PrE expansion culture conditions. To further quantify Oct4 expression, we generated *Oct4*^*T2A-EGFP*^ Scr and KO ESC lines (Fig. 3B). The presence of the T2A site leads to ribosomal skipping, resulted in translation of separate Oct4 and EGFP proteins. This facilitates focusing on transcriptional activity, as EGFP is not further coupled to the Oct4 post-transcriptional fate. Then we performed RA-mediated differentiation of the ESCs with daily EGFP monitoring throughout five-day period. EGFP signal in the Scr control cells was almost completely lost by the third day of differentiation, while nearly 37% of EGFP^+^ cells were observed by this time point in KO cells (Fig. 3C), which agrees with the above results (Fig. 3A). Further analysis revealed nearly 16% of EGFP^+^ KO cells by the fifth day of differentiation. To rule out that the EGFP^+^ cell population contains residual ESCs, we FACS-enriched this population on the day 4 of differentiation, washed these cells of RA, and cultured them in the SL medium for additional four days. Neither expansion of the EGFP^+^ cell population, nor re-emergence of morphologically distinct ESCs were detected (Suppl. Fig. 3A). We have also followed EGFP signal dynamics during differentiation proceeded via embryoid body formation (Suppl. Fig. 3B) or induced by LIF withdrawal (Suppl. Fig. 3C). In both differentiation scenarios, most of the *Pou5f1*^*T2A-EGFP*^ Scr cells lost EGFP signal by day 5, while a significant sub-population of the *Pou5f1*^*T2A-EGFP*^ KO cells retained EGFP expression. Therefore, all three examined types of directed ESC differentiation (RA-induced, induced by LIF withdrawal, or progressed via embryoid body formation) featured a delayed Oct4 expression decline is in the absence of Pcbp1 function.

**Figure 3.**
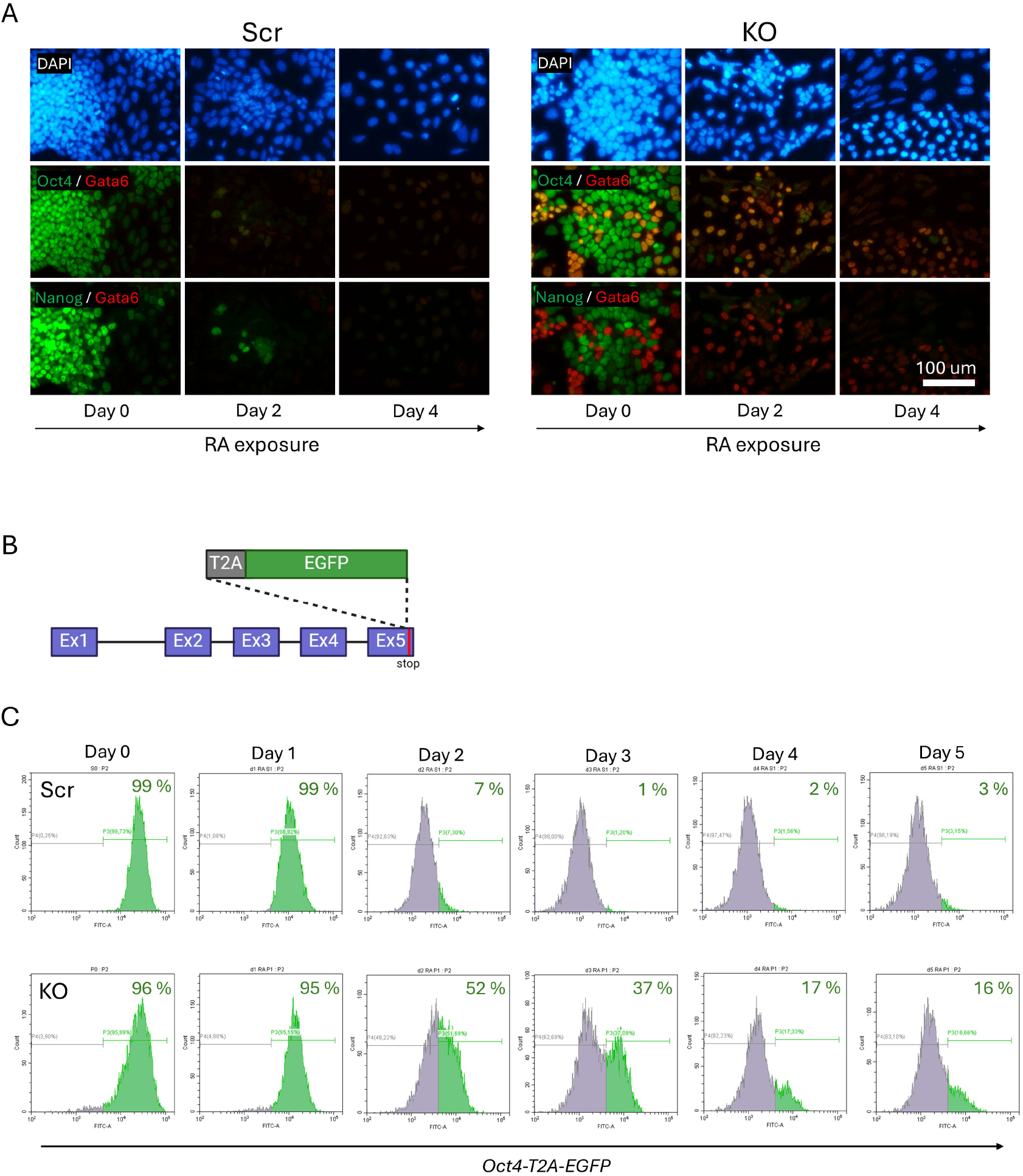
Oct4 down-regulation during RA-induced differentiation of KO ESCs is delayed. (A) Immunofluorescent staining of the Ser and KO ESCs for Oct4, Nanog, and Gata6 proteins during RA-induced differentiation at the indicated time points. (B) Schematic representation of the *T2A-EGFP* cassette inserted into the fifth Oct4 exon (Ex) just in front of the stop-codon. (C) Flow cytometry analysis of EGFP dynamics throughout the differentiation of the Oct4T2A*-EGFP* Ser and KO ESCs. Cells were cultured in the SL medium (Day 0) or in the same medium deprived of LIF and containing RA (Days 1-5).

### MEK-inhibition and Pcbp1-deficiency in PrE specification

It is known that PrE development is dependent on MEK/ERK-signaling [57]. Therefore, we have asked if the PrE cells spontaneously arising in KO ESC culture can be eliminated by the commonly used MEK-inhibitor PD0325901 (hereafter referred to as PD). We tested this inhibitor along with the GSK3β inhibitor CHIR99021 (CHIR), as these two small molecules are routinely used as components of the well-known 2i medium for the establishment of homogeneous ESC culture in the state of naïve pluripotency [58]. As expected, the addition of PD or both inhibitors, but not of CHIR alone, led to an elimination of Gata6^+^ PrE subpopulation in the KO ESC culture (Fig. 4A), which was accompanied by an enforced Nanog expression (Fig. 4B). Next, we asked if the exposure to PD restores normal Oct4 down-regulation during RA-mediated differentiation of the KO ESCs. Unexpectedly, however, the PD treatment applied prior to the differentiation significantly increased (up to 63%) the amount of residual EGFP^+^ KO cells by day 3 of RA-mediated differentiation (Fig. 4C, upper right panel), compared to 26% of the same cells non-treated with PD (Suppl. Fig. 4A). Also, the pre-treatment led to a delayed EGFP decline in differentiating Scr cells – 15% vs. 1% (Fig. 4C, upper left panel, and Suppl. Fig. 4A). Moreover, under the PD pre-treatment, Scr ESCs began to demonstrate PrE differentiation capacity (20% vs. 0.4%), revealed by Gata6 immunostaining with flow cytometry analysis (Fig. 4C, bottom left panel and Suppl. Fig. 4A). This result agrees with previously reported data showing that MEK inhibitor pre-treatment primes both mouse and human ESCs for PrE differentiation [59-61]. At the same time, population of Gata6^+^ PrE KO cells was enlarged to approximately 22% (Fig. 4C, bottom right panel), as compared to 10% of same cells non-primed with PD (Suppl. Fig. 4A). Immunocytochemical analysis shows Oct4/Gata6 colocalization in both Scr and KO cells (Suppl. Fig. 4B), implying Oct4 role in PrE specification. Summarizing these results, we can conclude that the ability to block spontaneous PrE differentiation of the KO ESCs by PD along with the observed additive effect of Pcbp1 knockout and PD pre-treatment on Oct4 down-regulation and enhanced PrE specification point to independent mechanisms of Pcpb1 and MEK/ERK-signaling action.

**Figure 4.**
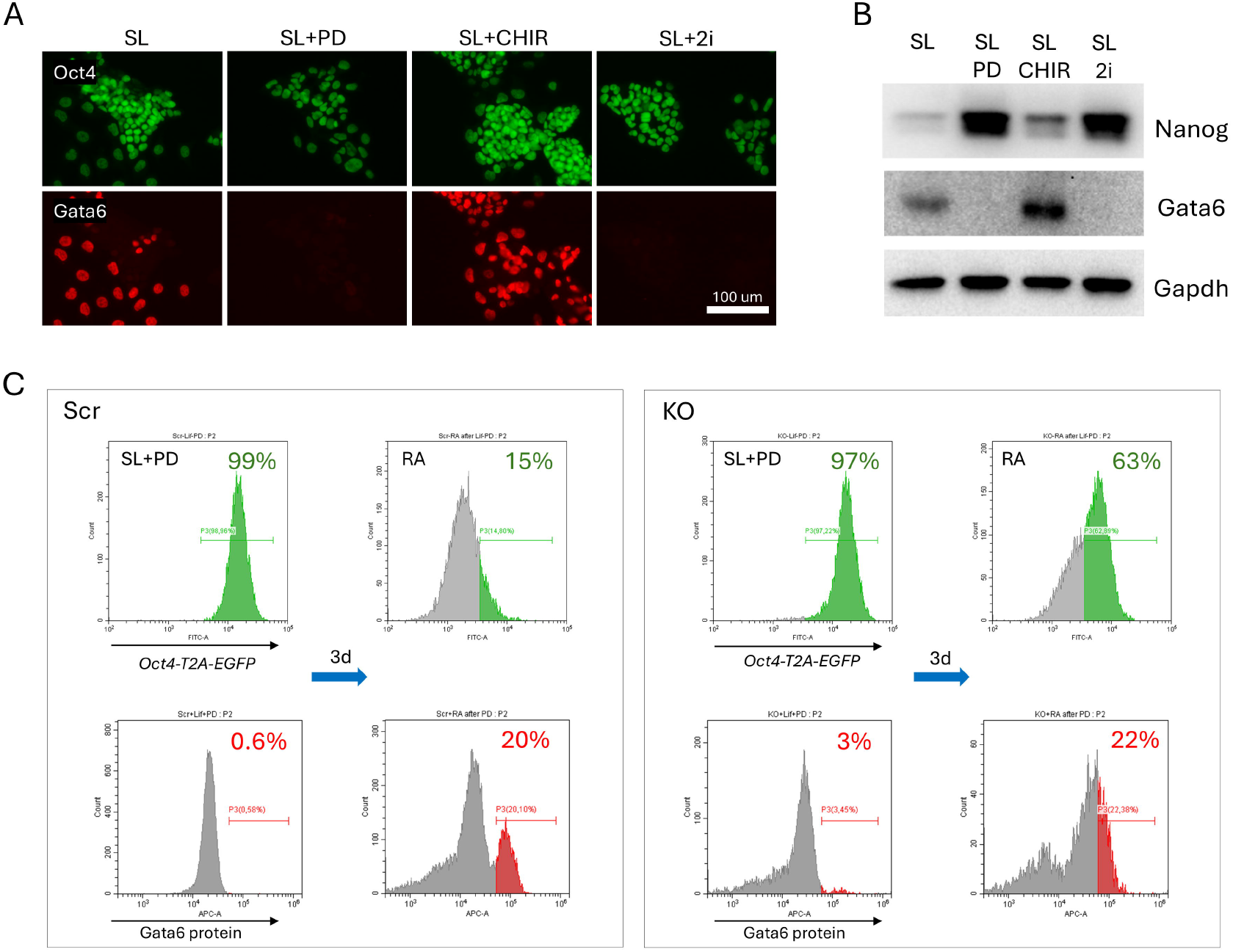
Dual role of MEK signaling in PrE specification. (A) Immunofluorescence staining for Oct4 and Gata6 proteins of the KO ESCs cultured in the SL medium supplemented with PD, CHIR, or both inhibitors (2i). (B) Western blot demonstrating the expression of Nanog and Gata6 in KO cells cultured as described in A. (C) Flow cytometry analysis of the Ser and KO ESCs pre-cultured in the SL supplemented with PD (SL+PD) and same cells additionally treated with RA for 3 days (without LIF and PD). Upper graphs and lower graphs represent the results of flow cytometry analysis analysis of the EGFP and Gata6 protein expression, respectively. Numbers represent the percentage of positive cells.

### Oct4 expression mediates Pcbp1-driven PrE differentiation

The above results indicate that Pcbp1 function loss may cause transcriptional dysregulation of *Pou5f1* gene, resulted in persistent Oct4 expression during pluripotency exit either via spontaneous or RA-triggered ESC differentiation, which, in turn, may promote the establishment of PrE specification program. To address this hypothesis, we made use of the mutant ESCs carrying conditionally targeted *Pou5f1* alleles, or *Pou5f1*^*flox/flox*^ [5] (Fig. 5A), along with 4-hydroxitamoxifen (4-OHT)-inducible ERT2-Cre-ERT2 recombinase knocked into a *Rosa26* allele, or *Rosa26*^*ERT2-Cre-ERT2*^. Addition of the 4-OHT to the culture medium leads to excision of the *Pou5f1* promoter and first exon (Suppl. Fig. 5A) with ensuing loss of Oct4 expression after 48 hours (Suppl. Fig. 5B). A successfully performed *Pcbp1* gene knockout in several sub-clones of one *Pou5f1*^*flox/flox*^;*Rosa26*^*ERT2-Cre-ERT2*^ ESC clone was confirmed by Western blotting (Suppl. Fig. 5C). To confirm that Oct4 expression mediates PrE differentiation driven by Pcbp1 loss and PD pre-treatment, we added 4-OHT one day before RA administration (Fig. 5B). In line with previous results, widespread Oct4^+^Gata6^+^ cell population emerged in control (- 4OHT) culture (Fig. 5C). In contrast, very few PrE cells were detected following 4-OHT treatment (Fig. 5D), and all these cells co-expressed Oct4, which is likely due to incomplete excision of the *Pou5f1*^*flox*^ alleles.

**Figure 5.**
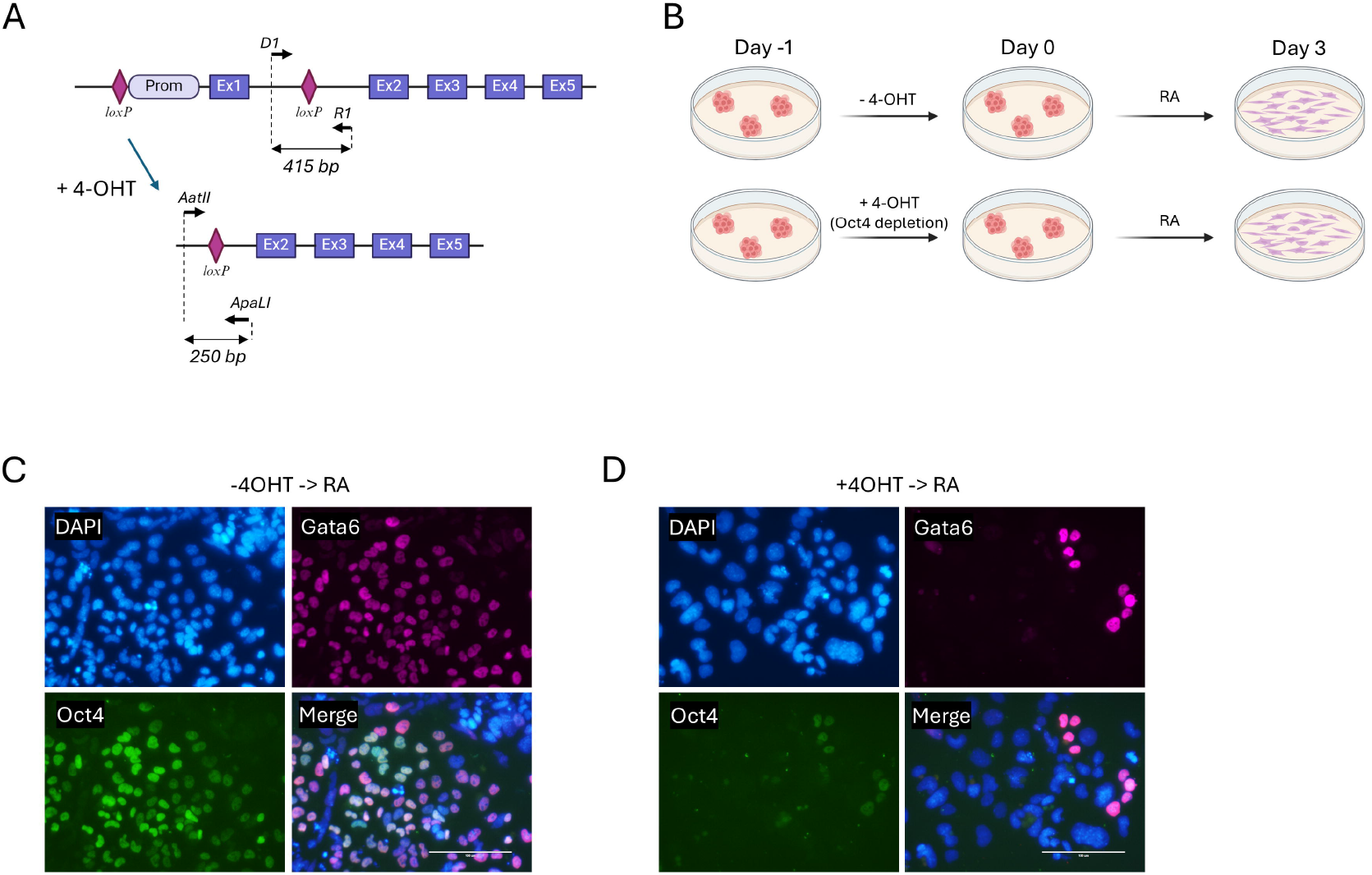
Oct4 expression mediates Pcbpl-driven PrE differentiation. (A) Schematic representation of the *Pou5fl* locus with promoter and first exon flanked by loxP sites, before and after 4-OHT exposure; locations of primers used for genotyping are indicated. (B) Schematic representation of strategy used for the Oct4 depletion via 4-OHT treatment in the KO “floxed” ESCs before RA-driven differentiation. (B) Scatter plots based on immunocytochemistry analysis showing Gata6 and Oct4 signal distribution in the KO ESCs without (left) or with (right) preliminary Oct4 depletion. (C, D) Immunocytochemical analysis of Gata6 and Oct4 expression in “floxed” ESCs without (C) or with (D) preliminary Oct4 depletion and differentiated by RA.

## Discussion

*Pou5f1* gene encoding the key pluripotency regulator Oct4 is subject to complex transcriptional and epigenetic regulation. Nevertheless, critical details about molecular mechanisms of this regulation are still missing. Multiple transcription factors, like Zscan4, Sall4, Nr5a2, Utf1, Tpt1, and Esrrb, have been found to bind *Pou5f1* regulatory elements *in vivo*, however, their loss in ESCs or pre-implantation embryos produced little or no effect on Oct4 expression [10, 23]. Same is true for hnRNP-K which binds the *Pou5f1*’s poly(C)-sites within the DE and PE but is dispensable for activation, maintenance, and repression of this gene [48]. It seems that *Pou5f1* has a flexible mechanism of activation with DE/PE being a part of super-enhancer [62]. We have proposed that the abovementioned transcription factors form multiprotein complexes on this and, perhaps, other super-enhancers involved in pluripotency control [62] and that individual components of this complex play highly redundant functions [48].

Like hnRNP-K, its close relative Pcbp1 seems to be involved in regulation of housekeeping function in undifferentiated ESCs. While the former protein is essential for ESC viability [48], the latter’s function is less crucial and might be related to cellular metabolism control or other general functions, as evidenced by the increase cell cycle length of the KO ESCs. Of note, relatively mild Pcbp1 phenotype in ESCs could be partly explained by a redundant function of a closely related homolog, Pcbp2, which is also expressed in ESCs. Importantly, unlike hnRNP-K, Pcbp1’s knockout results in slower proliferation and tendency to quit the pluripotent state. At the same time, it seems that PrE emergence during pluripotency shutdown is a consequence of sustained *Pou5f1* transcriptional activity. The latter idea is supported by the observations that Pcbp1 occupies *Pou5f1* DE and PE, while its loss delays Oct4 down-regulation during ESC differentiation. Besides, our data show that Oct4 depletion throughout KO ESC differentiation abolishes PrE program onset. This finding clarifies the mechanisms that are affected in KO cells and connects the observed *Pou5f1* dysregulation and PrE differentiation. Our study also substantiates the evidence that Oct4 out of LIF/STAT3 signaling directs PrE specification [63] and points to an essential role of Pcbp1 in synchronizing *Pou5f1* transcription with the pluripotency network.

We have demonstrated that MEK-inhibition prevents spontaneous PrE emergence of KO ESCs, but primes both control and KO cells for PrE during RA-mediated differentiation. Moreover, we observed an additive effect of Pcbp1 loss and PD treatment on both sustained Oct4 expression and PrE program onset during this differentiation, which indicates that Pcbp1 and MEK/ERK-signaling represent two different mechanisms driving PrE differentiation. It is known that MEK inhibition in early embryogenesis converts all ICM cells into pluripotent epiblast, while Fgf4 drives PrE differentiation [57, 64]. The MEK inhibitor PD is commonly used to push ESCs into naïve state, which resembles that of pre-implantation epiblast at E4.5 [58, 65, 66]. As ESCs under the SL conditions are closer to the primed pluripotent state than naïve ESCs [65], it seems that PD addition shifts ESC epigenetic landscape back to naïve state, which is more potent for PrE differentiation [59-61, 67].

There are still several open questions such as what molecular mechanism underlies the high rates of spontaneous differentiation of ESCs upon Pcbp1 function loss. There are rare spontaneous differentiation events under the SL conditions, but these events are greatly increased in number of KO ESCs. It is also notable, that artificially maintained Oct4 expression did not lead to a comparable PrE differentiation under the SL conditions and resulted in PrE differentiation only after RA treatment. Lentiviral infection applied to rescue Oct4-deficient ESCs results in random insertion of the construct throughout the genome. Due to extensive genome methylation during differentiation [68, 69], some lentivirus copies are unavoidably subject to silencing, resulted in Oct4 level decrease and reduced yield of PrE cells. Future attempts with *Rosa26*-intergated inducible *Pou5f1* transgenes may clarify this issue. Very recent study has shown that the Oct4^+^ PrE cells are still able to de-differentiate and contribute to trophectoderm, pluripotent epiblast, and primitive endoderm both *in vitro* and during embryogenesis [70]. Our KO PrE cells are characterized by a sustained Oct4 expression and thus, potentially, feature two-way ESC-PrE transitions. At last, the observed continued Oct4 expression in KO cells during differentiation assumes that some other factors drive its transcription in the absence of Pcbp1. There is an intriguing possibility that there are some other members of poly(C)-binding transcription factor families, which compete with Pcbp1 for binding to the *Pou5f1* regulatory elements. Future studies may shed light on these issues, as well as extrapolate the presented results on early embryogenesis.

## Author contributions

E.I.B. conceived the study, designed experiments, performed major part of experimental work and wrote the first draft; A.S.Z. contributed to western blot, immunocytochemistry and genotyping; M.N.G. performed *EGFP* gene integration into the *Pou5f1* locus in Scr and KO cells; A.S.Z., M.N.G., E.E.P. contributed to cell culture work; A.A.K. modified *Pou5f1*^*flox/flox*^ ESCs by integrating *Cre*-recombinase gene into the *ROSA26* locus and established plasmids for CRISPR/Cas9-mediated *EGFP* gene insertion into the *Pou5f1* locus; D.V.S. performed RT-PCR analyzes and contributed to figures design; N.D.A. performed flow cytometry analysis and FACS; A.N.T. conceived the study, designed experiments, contributed materials, tools and reagents, edited and approved final version of the manuscript.

## Acknowledgments

The study was supported by the Russian Science Foundation (RSF) grant № 23-75-10096, https://rscf.ru/en/project/23-75-10096/. Schematic images were created with BioRender.com.

