## Supplementary figures for "Pcbp1 constrains Oct4 expression in the context of pluripotency"

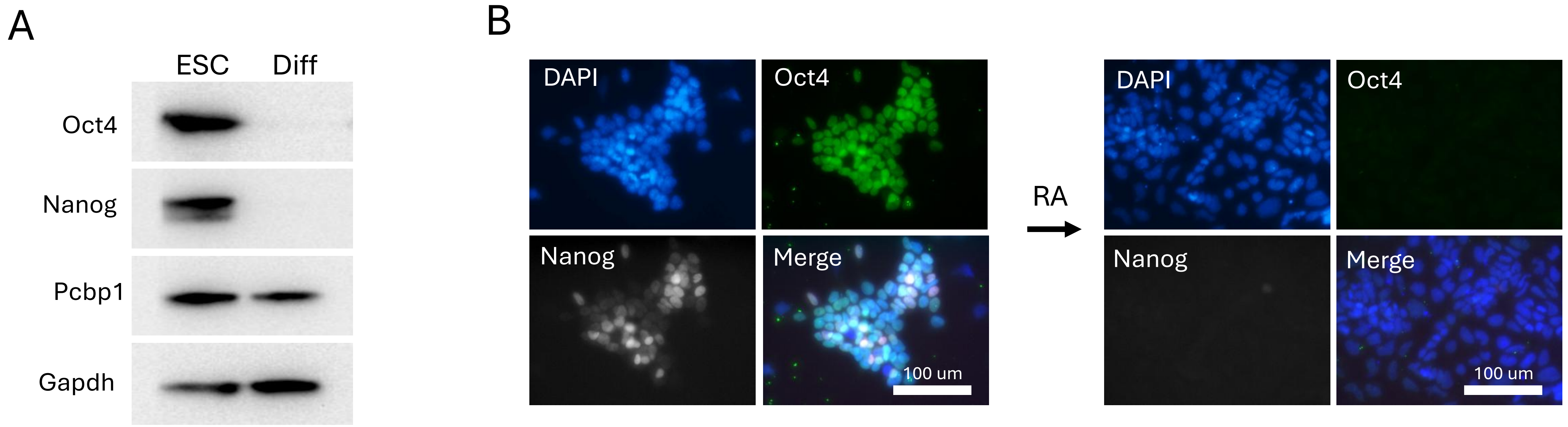


**Supplementary Figure 1. Verification of RA-mediated ESC differentiation.** (A) Western blot showing Oct4, Nanog, and Pcbp1 expression in ESCs cultured in the SL medium and after three days after the addition of RA and LIF withdrawal (Diff); (B) Immunofluorescence staining for Oct4 and Nanog in ESCs following 3-day differentiation induced with RA.

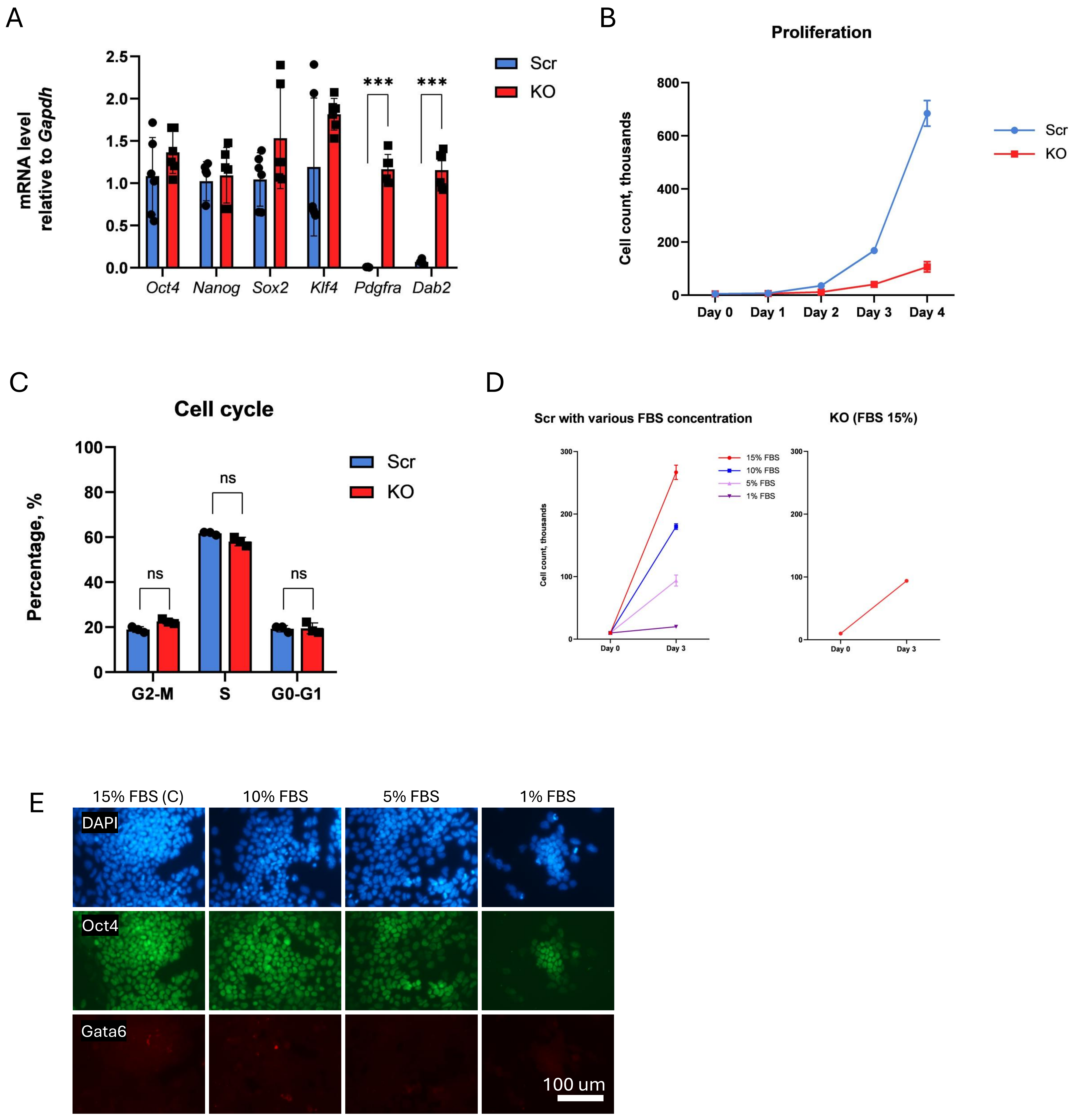


**Supplementary Figure 2. Characterization of the KO ESCs.** (A) RT-PCR analysis of pluripotency-related (Oct4, Nanog, Sox2, Klf4) and PrE-specific marker expression (Pdgfra and Dab2) in the Scr and KO independently derived ESC clones. Groups were compared using the Mann–Whitney non-parametric test (N=3, n=2). (B) Proliferation analysis of Scr and KO ESCs cultured under the SL conditions (N=3). (C) Flow cytometry analysis of cell cycle phase distribution after DAPI staining of the Scr and KO ESC clones (N=3). (D) Analysis of Scr ESC proliferation in the SL medium containing indicated concentrations FBS (left panel). Proliferation kinetics of the KO ESCs in the standard SL medium containing 15% of FBS (right panel) (N=3). (E) Immunofluorescence staining of Oct4 and Gata6 in the Scr ESCs cultured for one week in the SL medium containing indicated concentrations of FBS.

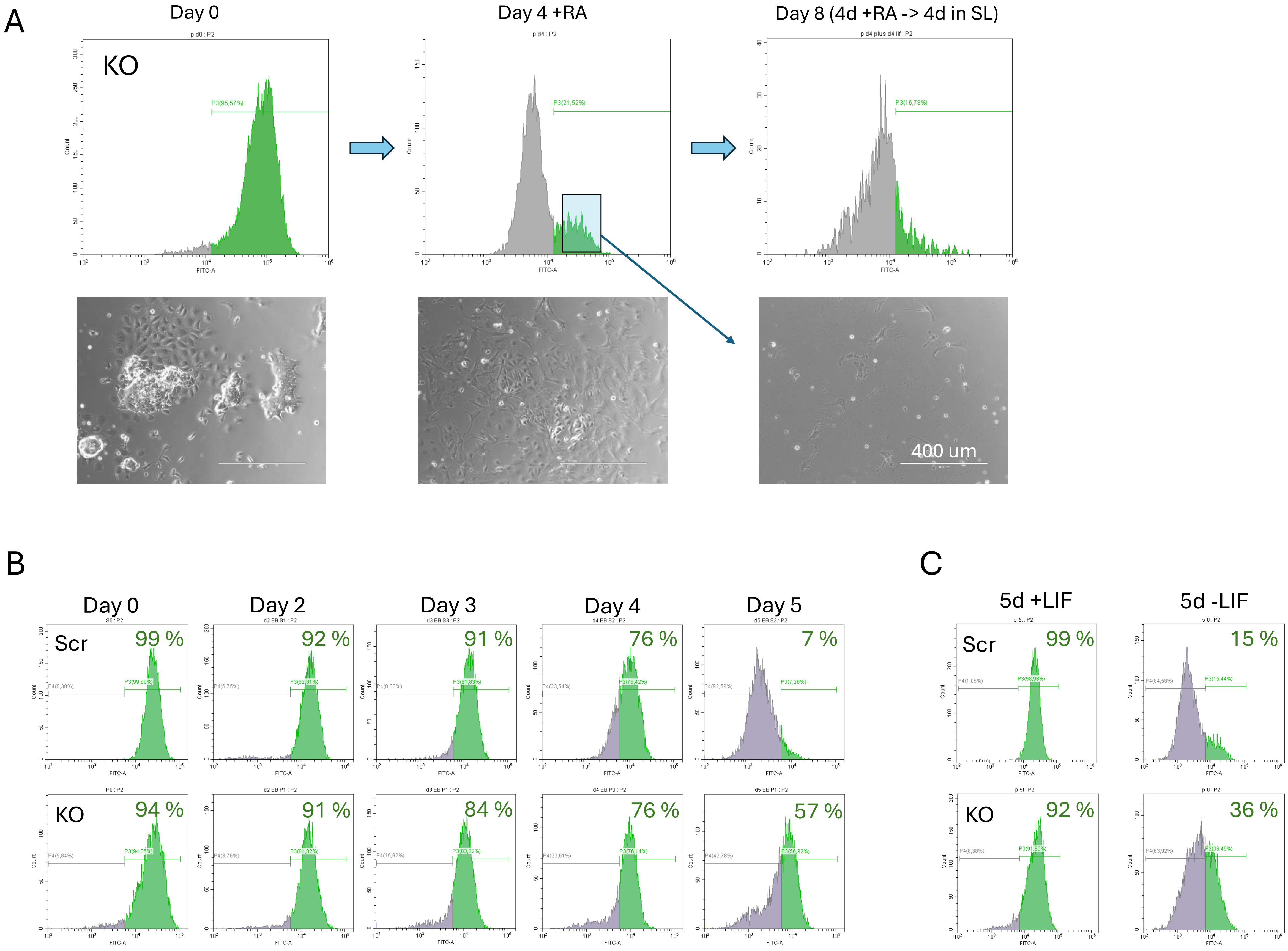


**Supplementary Figure 3. EGFP down-regulation dynamics during differentiation of *Oct4*$^{T2A\text{-}EGFP}$ Scr and KO ESCs.** (A) Flow cytometry analysis of EGFP fluorescence in the *Oct4*$^{T2A\text{-}EGFP}$ KO cells (upper panels) and corresponding phase contrast microphotographs of these cells (lower panels) cultured in the SL (left panels); the cells were then cultured for 4 days in medium containing RA (middle panel); EGFP$^{+}$ were isolated and cultured for additional 4 days in the SL (right panels. (B) Flow cytometry analysis of EGFP expression (shown as percentage of EGFP$^{+}$ cells) in the *Oct4*$^{T2A\text{-}EGFP}$ Scr and KO cell cultures at different time points of differentiation in embryoid bodies (Days 0-5). (C) Same analysis performed after 5-day culturing in the presence or absence of LIF, as indicated.

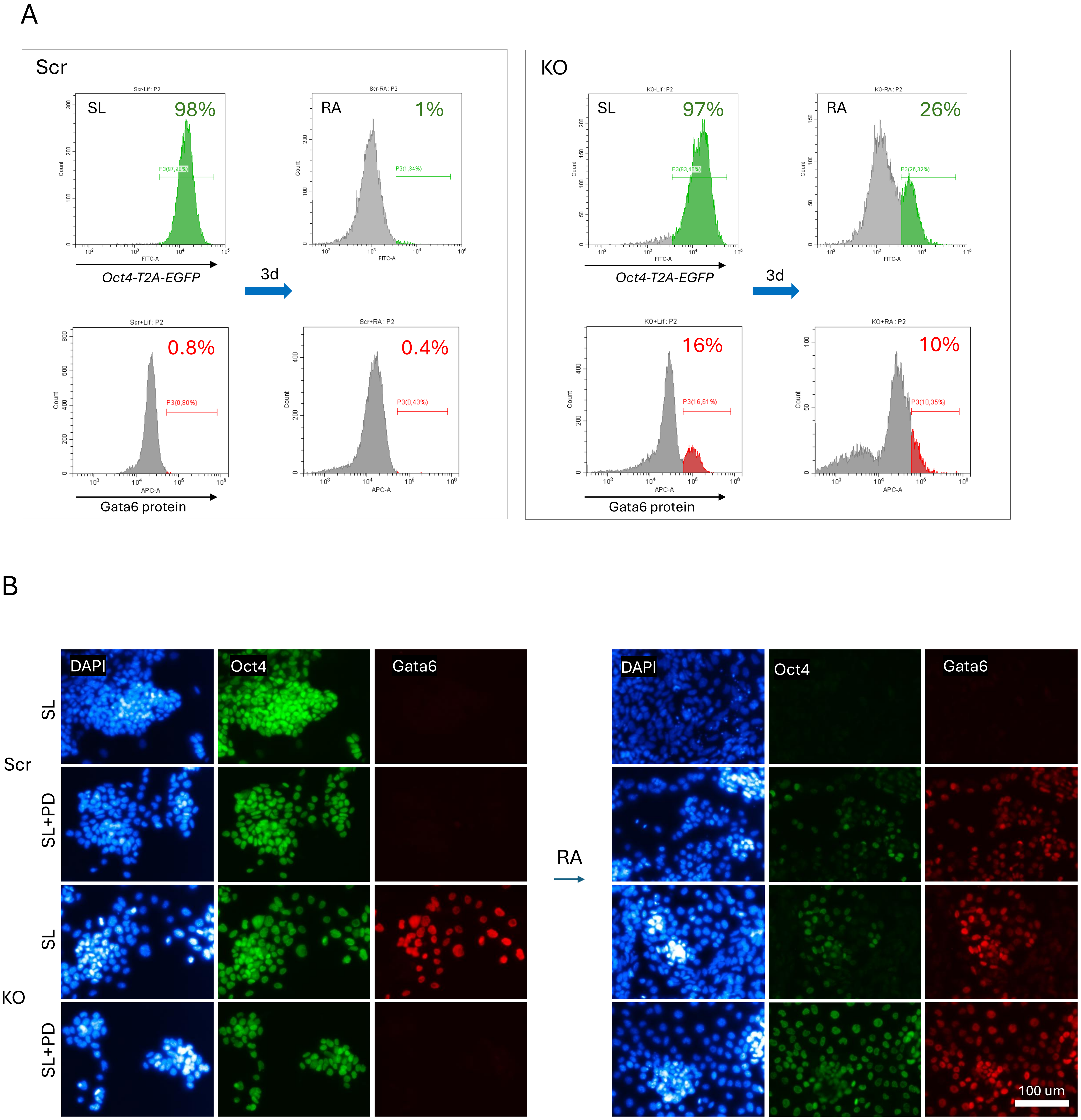


**Supplementary Figure 4. MEK-inhibition primes ESCs for PrE differentiation**. (A) Reference flow cytometry analysis of the Scr and KO ESCs pre-cultured in the SL (i.e. without the addition of PD) and same cells additionally treated with RA for 3 days (without LIF). Upper graphs and lower graphs represent the results of flow cytometry analysis analysis of EGFP and Gata6 protein expression, respectively. Numbers represent percentage of corresponding cells. (B) Immunofluorescent analysis of Oct4 and Gata6 expression in the Scr and KO ESCs cultured in the SL or SL+PD media and differentiated for three days in the presence of RA (without LIF and PD).

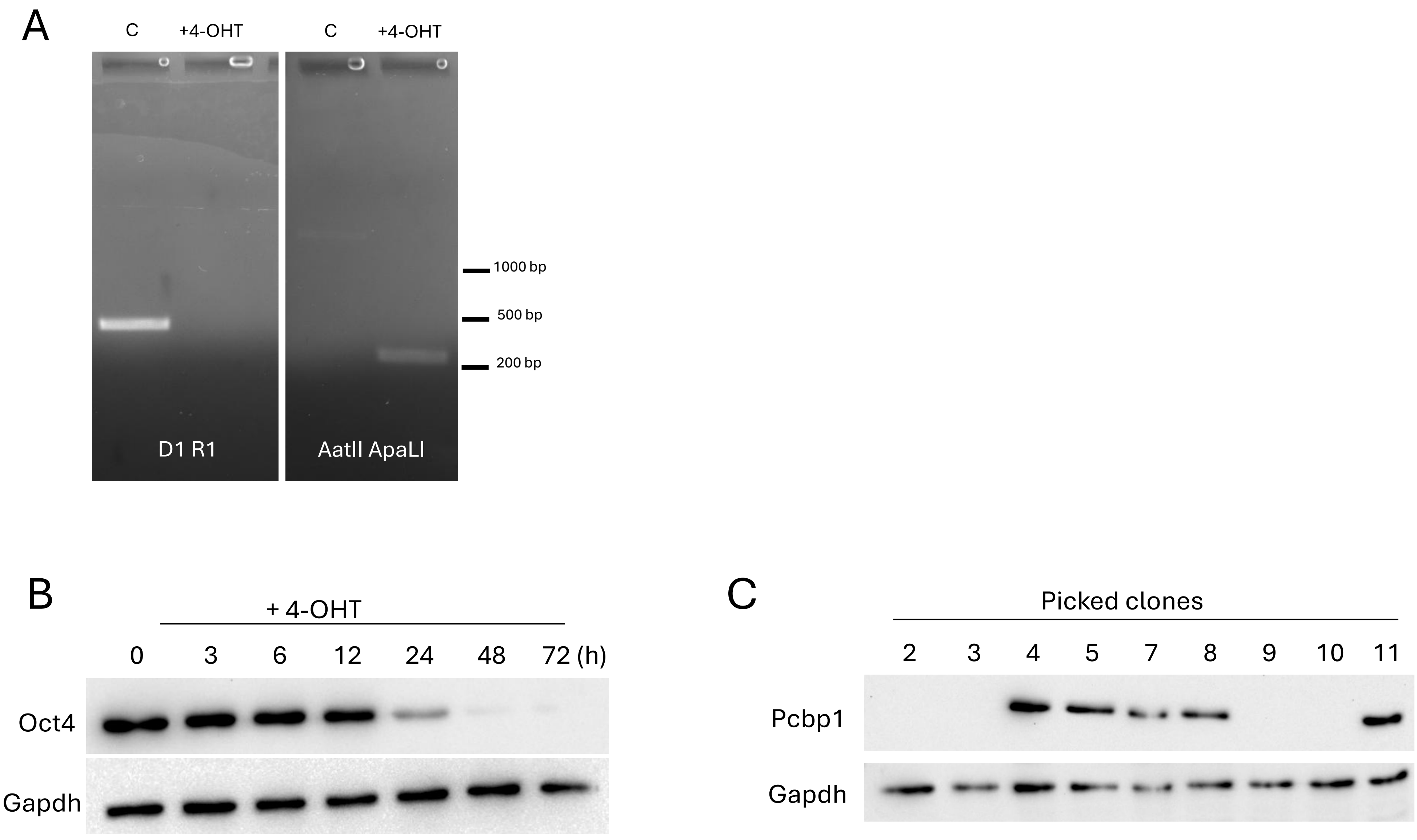


**Supplementary Figure 5. *Pcbp1* knockout performance in "floxed" ESCs.** (A) PCR-genotyping of the "floxed" ESCs before and after 4-OHT administration. (B) Oct4 expression in "floxed" ESCs at different timepoints after 4-OHT administration. (C) Pcbp1 expression check after CRISPR/Cas9-mediated *Pcbp1* knockout.
